# Rigor and Transparency Index, a new metric of quality for assessing biological and medical science methods

**DOI:** 10.1101/2020.01.15.908111

**Authors:** Joe Menke, Martijn Roelandse, Burak Ozyurt, Maryann Martone, Anita Bandrowski

## Abstract

The reproducibility crisis in science is a multifaceted problem involving practices and incentives, both in the laboratory and in publication. Fortunately, some of the root causes are known and can be addressed by scientists and authors alike. After careful consideration of the available literature, the National Institutes of Health identified several key problems with the way that scientists conduct and report their research and introduced guidelines to improve the rigor and reproducibility of pre-clinical studies. Many journals have implemented policies addressing these same criteria. We currently have, however, no comprehensive data on how these guidelines are impacting the reporting of research. Using SciScore, an automated tool developed to review the methods sections of manuscripts for the presence of criteria associated with the NIH and other reporting guidelines, e.g., ARRIVE, RRIDs, we have analyzed ~1.6 million PubMed Central papers to determine the degree to which articles were addressing these criteria. The tool scores each paper on a ten point scale identifying sentences that are associated with compliance with criteria associated with increased rigor (5 pts) and those associated with key resource identification and authentication (5 pts). From these data, we have built the Rigor and Transparency Index, which is the average score for analyzed papers in a particular journal. Our analyses show that the average score over all journals has increased since 1997, but remains below five, indicating that less than half of the rigor and reproducibility criteria are routinely addressed by authors. To analyze the data further, we examined the prevalence of individual criteria across the literature, e.g., the reporting of a subject’s sex (21-37% of studies between 1997 and 2019), the inclusion of sample size calculations (2-10%), whether the study addressed blinding (3-9%), or the identifiability of key biological resources such as antibodies (11-43%), transgenic organisms (14-22%), and cell lines (33-39%). The greatest increase in prevalence for rigor criteria was seen in the use of randomization of subjects (10-30%), while software tool identifiability improved the most among key resource types (42-87%). We further analyzed individual journals over time that had implemented specific author guidelines covering rigor criteria, and found that in some journals, they had a big impact, whereas in others they did not. We speculate that unless they are enforced, author guidelines alone do little to improve the number of criteria addressed by authors. Our Rigor and Transparency Index did not correlate with the impact factors of journals.

## Introduction

The National Institutes of Health (NIH) have designed and adopted a set of rigor and reproducibility guidelines expected to be addressed in grant proposals submitted to the NIH that covers the aspects of study design most likely to impact a study’s reproducibility (for NIH Guidelines see NOT-OD-15-103^1^; See also EU Report Open Science Monitoring^2^; for their intellectual underpinning^3^; for examples^4,5^). Multiple journals have adopted similar guidelines in their instructions to authors (e.g. Nature Checklist^6^).

The NIH guidelines are part of a growing list of recommendations and requirements designed to address different aspects of rigor and reproducibility in the scientific literature, e.g., the ARRIVE^7^, CONSORT^8^ and RRID^9^ standards. The Animal Research: Reporting of *In Vivo* Experiments (ARRIVE) guidelines are a highly comprehensive and universally accepted set of criteria that should be addressed in every animal-based experiment. The guideline contains 39 items (20 primary questions and 19 subquestions). The Consolidated Standards of Reporting Trials (CONSORT) statement is comprised of a 25 item checklist along with a flow diagram governing how clinical trials should be reported. The RRID Initiative, another reproducibility improvement strategy, asks authors to add persistent unique identifiers called research resource identifiers (RRIDs) to disambiguate specific assets used during experimentation. RRIDs can be considered as universal product codes (UPC) that identify the ingredients needed for an experiment. The initiative covers a wide variety of resources including (but not limited to): antibodies, plasmids, cell lines, model organisms, and software tools. The initiative was started because antibodies were notoriously difficult to identify unambiguously in the published literature.^10^

Unfortunately, studies of publishing practices generally find poor compliance by authors and enforcement by reviewers, even with the availability of checklists and instructions to authors. Even when authors assert that they follow ARRIVE, the evidence still shows that the guidelines are not followed.^11,12^ In the case of RRIDs, the guidelines were not routinely followed when authors were asked by journals through instructions to authors or by checklists, but a direct request from the editors for RRIDs during the publication process proved highly effective in improving author compliance.^9^

In order to reduce confusion around the proliferation of guidelines and to improve author compliance, the above guidelines were incorporated into the Materials Design, Analysis, and Reporting (MDAR) framework, a recently released pan-publisher initiative enacted to create a consistent, minimum reporting standard that spans across all life sciences.^13^ The MDAR checklist includes many of the elements that are present in the NIH, ARRIVE, CONSORT and RRID standards. The checklist is not intended to supplant more granular reporting of information, but rather is to be used as a generalist instrument across the biological research community.

One reason that author compliance may be low is not simply that there are a plethora of guidelines, but rather that there are few tools that support authors, reviewers and editors in implementing and enforcing such guidelines. Similarly, studies that seek to investigate the degree to which authors comply have been limited to manual review of a few criteria.^14,15^ We recently developed SciScore, an automated tool using natural language processing (NLP) and machine learning, that can be used by journals and authors to aid in compliance with the above guidelines. SciScore currently evaluates compliance with six key recommendations from the above guidelines and checks for key resource identifiability for a variety of resource types. Here we introduce SciScore and show how it can be used to assess the impact of rigor and reproducibility reporting guidelines comprehensively across the open access scientific literature. Using SciScore, it is now possible to create a Rigor and Transparency Index across journals that can be compared to current metrics such as Impact Factor.

## Methods and Materials

### Text mining the open access subset of PubMed Central

For this study, we downloaded and processed all open access literature available through PubMed Central (PMC, RRID:SCR_004166) in September of 2019. In total, we obtained data from 1,578,964 articles from 4,686 unique journals. The PMC Open Archives Initiative (OAI) dataset was initially downloaded as directories (one per journal named by the journal’s standard abbreviation) allowing for a clear differentiation of each journal. Articles only available as PDFs were not included in the PMC-OAI dataset, and were therefore excluded from our analysis. In addition, abstract-only articles and articles without a methods section were also excluded from our analysis because the reporting criteria are generally included only in the materials and methods. We limited our analysis to journals that had published more than 10 papers for any given year in the PMC-OAI.

In order to create the dataset used for our analysis, the OAI articles were fed through the SciScore™ named-entity recognition classifiers. SciScore currently uses 6 core named-entity recognition classifiers (see Table 1 for a complete list of entity types detected). Each of these was validated using precision and recall as well as their harmonic mean, F1. The values for each entity type are listed in Table 1.

**Table 1.** Individual Classifier Performance for Named-Entities

| Entity Type | F1 | Precision | Recall | Training Set Size<br>(sentences<br>w/entities) |
| --- | --- | --- | --- | --- |
|  | Mean ± SD | Mean ± SD | Mean ± SD |  |
| Rigor Criteria (5 total points) |  |  |  |  |
| Institutional Review Board Statement * | 81.41 ± 3.62 | 84.45 ± 5.26 | 79.57 ± 8.83 | 340 |
| Consent Statement * | 94.75 ± 1.68 | 96.29 ± 2.42 | 93.38 ± 3.63 | 373 |
| Institutional Animal Care and Use<br>Committee Statement * | 81.30 ± 4.20 | 89.30 ± 4.60 | 74.89 ± 6.12 | 591 |
| Randomization of subjects into groups | 83.05 ± 3.04 | 80.25 ± 5.05 | 86.45 ± 4.64 | 368 |
| Blinding of investigator or analysis | 78.96 ± 12.38 | 77.74 ± 17.16 | 81.79 ± 10.32 | 183 |
| Power analysis for group size | 64.45 ± 29.37 | 73.74 ± 34.13 | 59.50 ± 26.91 | 81 |
| Sex as a biological variable | 88.32 ± 3.91 | 87.94 ± 6.03 | 88.93 ± 3.52 | 862 |
| Cell Line Authentication ^ | 54.08 ± 11.88 | 85.70 ± 10.78 | 41.15 ± 12.82 | 155 |
| Cell Line Contamination Check ^ | 91.70 ± 5.24 | 93.35 ± 7.15 | 90.65 ± 7.05 | 151 |
| Key Biological Resources (5 total points) |  |  |  |  |
| Antibody | 78.94 ± 2.62 | 86.89 ± 3.78 | 72.46 ± 3.20 | 16,772 |
| Organism | 66.05 ± 4.70 | 79.91 ± 6.28 | 56.64 ± 5.75 | 4,439 |
| Cell Line | 70.07 ± 5.95 | 86.48 ± 3.27 | 59.34 ± 8.03 | 1,763 |
| Plasmid ** | 79.62 ± 3.35 | 92.53 ± 3.80 | 70.09 ± 4.85 | 2,568 |
| Oligonucleotide ** | 83.03 ± 9.05 | 95.28 ± 3.13 | 74.94 ± 13.90 | 1,893 |
| Software Project/Tool | 89.03 ± 0.90 | 92.49 ± 2.08 | 85.84 ± 1.10 | 10,161 |

\* Institutional review board and consent statements are scored together as a block where detection of one or more of these entities will give the full point value for this section.
^ Cell line authentication and contamination statements are only scored when a cell line is detected in the key resources table and they are scored together, either of these will provide the full points for this section.
\*\* Entity type not used for analysis in the current paper.

Each classifier component was trained and tested separately for precision and recall using human curated data. The curator labeled each entity type within tens of thousands of example sentences using the smallest word chunk that was still informative. For example, in the sentence, “We used the mouse monoclonal GFAP antibody from Sigma”, the antibody name would be [GFAP], while the clonality would be [monoclonal]. The source organism would be [mouse], and the antibody vendor would be [Sigma]. For antibody detection, these and any other individual components describing antibodies are used to identify an antibody and its additional metadata. We assume that the authors will report some, but not all antibody features for any given antibody. Treating the antibody name feature as its primary tag, the overall F1 score for antibodies is 78.9 with a precision of 86.9 and a recall of 72.5. If the curator and algorithm did not have complete agreement with regard to the entity in our training dataset, it was considered a miss e.g., [GFAP] vs [GFAP antibody] in the example sentence. The remaining features were used to improve our antibody name recognition and to determine whether the antibody was identifiable. We tested these values using 10 random splits of the data where 90% of the human curated data was used as training and 10% is used as the test. The final value comes from a mean of all 10 training trials. If the F1 was determined to be below the desired 70% threshold for key resources, we attempted to increase the training dataset size. We did not set a minimum desired F1 threshold for our rigor classifiers as training data was far more difficult to locate for certain criteria. Overall, 11 curators have worked on annotating various components over the last 4 years.

While the cell line algorithm has been tested previously to find the total number of cell lines used throughout the open access subset of PubMed Central,^16^ the other algorithms had not been thoroughly validated before this on complete articles outside of the training set. To validate SciScore’s total performance, we tested SciScore against an independent set of human-curated data. This set was created using 250 papers randomly chosen using the random() function in SQLite (RRID:SCR_017672) from our dataset of open-access papers. Each paper was then manually reviewed by a curator to determine which rigor criteria and key resource information had been referenced. For each paper, the methods section was read, and the curator noted the presence or absence of each entity type. For this check, the curator and SciScore were considered to be in agreement if both had marked an entity type as either present or absent. If there was a disagreement, it would then be classified as either a false negative error or a false positive error. For example, if a paper containing multiple antibodies was noted by the curator as having antibodies present, and SciScore determined that there were antibodies present as well, then this would be considered an agreement. In that example, if SciScore had determined that no antibodies were present, then this would be considered a false negative error. Note, the curator did not keep track of exactly which antibodies were used in the paper or how many. For this analysis, the curator was blinded to the output of the algorithm while curating papers in this set. For validation, this information was then compared against our calculated SciScore classifier performances, listed in Table 1; the results of this analysis are in Table 2.

**Table 2:** Rates of false negatives, false positives, and overall agreement based on manual analysis of 250 scored papers (SciScore > 0) from our dataset.

| Entity Type | False Positives |  | False Negatives |  | Overall Agreement |  |
| --- | --- | --- | --- | --- | --- | --- |
|  | Size (#) | Rate (%) | Size (#) | Rate (%) | Size (# agreed) | Rate (%) |
| **<u>Rigor Criteria</u>** |  |  |  |  |  |  |
| Institutional Review Board Statement or Consent Statement | 11 | 4.4 | 3 | 1.2 | 236 | 94.4 |
| Institutional Animal Care and Use Committee Statement | 7 | 2.8 | 15 | 6.0 | 228 | 91.2 |
| Randomization of subjects into groups | 16 | 6.4 | 8 | 3.2 | 226 | 90.4 |
| Blinding of investigator or analysis | 2 | 0.8 | 7 | 2.8 | 241 | 96.4 |
| Power analysis for group size | 12 | 4.8 | 6 | 2.4 | 232 | 92.8 |
| Sex as a biological variable | 5 | 2.0 | 20 | 8.0 | 225 | 90.0 |
| Cell Line Authentication or Contamination Check | 12 | 4.8 | 0 | 0.0 | 238 | 95.2 |
| **<u>Key Biological Resources</u>** |  |  |  |  |  |  |
| Antibody | 2 | 0.8 | 3 | 1.2 | 245 | 98.0 |
| Organism | 3 | 1.2 | 7 | 2.8 | 240 | 96.0 |
| Cell Line | 6 | 2.4 | 4 | 1.6 | 240 | 96.0 |
| Software Project/Tool | 8 | 3.2 | 41 | 16.4 | 201 | 80.4 |

### Scoring Framework

All research papers in the PMC-OAI dataset were scored on a 10-point scale. To calculate the total score for each paper, the scoring was broken down into two equally weighted sections: 5 points for rigor adherence (made up of the rigor criteria listed in Table 1) and 5 points for key resource identification (consisting of the key biological resource types listed in Table 1). In cases where no rigor criteria or key resources were detected, the paper received a score of 0. Papers given a 0 were excluded from the dataset because in cases where SciScore cannot find any criteria to judge, there is no way of determining if a score is appropriate. As SciScore was originally intended for biomedical research articles, papers scored as 0 typically fall far outside of its scope (e.g. X-ray crystallography), or are the wrong paper type (e.g. a letter to the editor). Indeed, of the 197,892 not applicable (0 scoring) papers, over 30,000 came from the following five journals: Acta crystallographica. Section E, Structure reports online (98% of articles scoring 0), Nanoscale research letters (71%), Beilstein journal of organic chemistry (78%), Acta crystallographica. Section E, Crystallographic communications (95%), and iScience (100%); see supplementary data file 1. In order to validate this assumption, a second set of human-curated data was created using 250 papers that had received a score of 0. These papers were randomly chosen using the random() function in SQLite. Each paper was then manually reviewed by a curator to determine if any rigor criteria had been mentioned and which key resources, if any, had been referenced. Similar to our scored paper analysis, any criteria found was marked as either present or absent. The curator was not blinded to the output of the algorithm for this set, which may introduce an element of bias for this portion of the analysis. We note that when creating the manually checked datasets, we grouped IRB and consent as well as cell line authentication and contamination statements so the coding would be consistent with the output of the automated pipeline.

The rigor section of each score adds roughly one point for each criterion that is detected. For example, the score would be increased by one point if the author discussed blinding of the investigator during data analysis. In certain cases, a particular criterion may be deemed irrelevant and is not scored, such as the cell line authentication statement, which would not be required in papers that do not use cell lines. Currently, we weigh ethical approval sentences (which could be of the following types: IRB, IACUC, or consent statement) as two points because this tool is intended for manuscripts in preparation and not having a statement on ethics can have serious legal ramifications. In short, simply comparing the total number of found, relevant criteria to the total number of expected, relevant criteria roughly provides the score for the rigor section. This presents a positive bias in scores towards vertebrate animal and human subjects papers that include the ethical approval statement, and a negative bias against cell line and invertebrate papers, as ethical approval is not required in those cases. The current version of the tool does not score cell line authentication if no cell line is detected, but does not yet have the ability to distinguish whether ethical approval is necessary.

The key resources section is scored altogether as one block and takes into consideration the total number of resources found using a similar found:expected ratio scoring system. In brief, all detected resources are categorized into two scoring groups: detected but not uniquely identifiable (no points), and identifiable (full points). In our example mentioned above, “We used the mouse monoclonal GFAP antibody from Sigma”, the algorithm is likely to detect a single antibody and vendor, but the catalog number or research resource identifier (RRID) would not be found. For this sentence, this resource would not contribute any points towards the “found” total because the resource is not uniquely identifiable. It would, however, still contribute towards the expected resources count, so if this was the only resource detected, the author would receive a 0 of 5 for this section. If the author were to provide a catalog number, the algorithm may suggest a RRID given that it is able to estimate with a high level of confidence a single RRID entity with matching metadata (this is granted a point for for identifiable section), a valid matching RRID guarantees the point. We then calculate the key resource section’s score using a weighted average based on these two categories: unidentifiable and identifiable, and the proportions of key resources in each. When the algorithm fails to recognize a resource, that is considered a false negative, occuring at rates outlined in Table 1. We note that the values reported in Table 1 are for individual entities, when an entity is discussed several times, the probability should be additive. Papers tend to discuss resources several times in the methods section; for cell lines, each cell line was mentioned twice, improving the rate of resource identification in the paper. So we expect that our scores for SciScore to curator agreement should be at or above the raw values. Final scores are rounded to the nearest integer.

### Impact Factor Comparison

All journals contained in the PMC-OAI were initially considered for our analysis. In order to ensure that the average score calculated for each journal was representative of their true average, we limited our analysis to journals with sample sizes larger than the minimum required sample size calculated for each journal. Journals that did not meet this minimum were excluded from our analysis. We then searched the Journal Citation Reports (JCR, RRID:SCR_017656) database (operated by Clarivate Analytics) to obtain the journal impact factor (JIF) and average JIF percentile for each journal’s 2018 scores. These values are the most recent obtainable scores as new JIF information is usually released ~6 months after the end of the year (e.g. JIF values for 2019 will be released around June of 2020). Searches were performed in November of 2019. Journals that did not have their information listed in the JCR were excluded from our analysis. JIF is “calculated by dividing the number of current year citations to the source items published in that journal during the previous two years” according to Clarivate Analytics (RRID:SCR_017657), the official source for JIFs. For JIFs in 2018, this roughly translates to the following equation (Eq. 1):

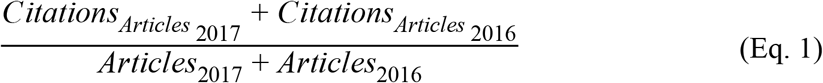

Because of this, when we calculated the average score for each journal, we only included scores from 2016 and 2017, so that the SciScores and JIFs would theoretically be representative of roughly the same papers. We say “roughly” because JIF is calculated using “citable items”, a vague term sometimes made up of a variety of article types (original research, commentaries, opinions, etc.),^17^ while SciScore is currently intended for use on original research. The average JIF percentile is calculated using the rank of each journal’s impact factor grouped by the field in which the journal is indexed. This calculation accounts for citation variations between different scientific fields as JIF percentile only compares journals within a specific category (cell biology journal vs. cell biology journal). As a result, any difference in citation counts between fields (e.g. physical chemistry vs. immunology) will be mitigated, allowing for a better comparison across all biomedical research. SciScore percentile was calculated based on the average SciScore of all 490 journals used in our impact factor comparison. In order to evaluate the correlations between JIF vs. SciScore and JIF percentile vs. SciScore percentile, we used Google Sheets to calculate Spearman’s rank-order correlation for each. Spearman’s correlation was chosen over Pearson’s because we did not assume bivariate normality. One potential source of bias affecting this analysis is the FUTON (full text on the Net) bias, which positively impacts citation counts for open-access research, while negatively impacting the number of citations for research not freely available on the web.^18,19^

### Statistics

To determine if a journal sampling was representative of its population in our impact factor analysis, we calculated the minimum sample sizes (n) required for each journal using the following equation (Eq. 2) for the sample size estimation of a finite population:

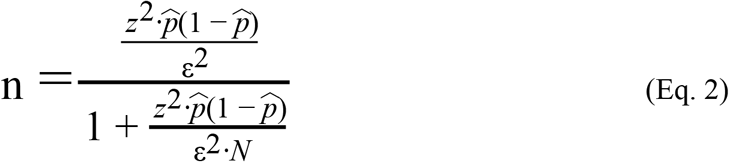

where z is the z score, 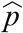 is the sample proportion, ε is the confidence interval, and N is the population. We used a confidence level of 95%, a confidence interval of 5%, and a sample proportion of ~0.875, which was the proportion of papers in our dataset that received a score above 0. Population sizes varied, but were determined by performing searches on PubMed restricted by publication type [journal article] and journal name. The minimum sample size was also calculated for each year to determine how far back our analysis should consider. For those calculations, the population was determined by the number of journal articles published in PubMed for a given year. These calculations were performed in Sheets (Google Sheets; RRID:SCR_017679). For all other analyses, journals were only included if more than 10 papers were scored per year.

For SciScore named-entity classifiers, we used the standard measures used to quantify performance: recall (R), precision (P), and the harmonic mean of R and P (F1). These were determined by the following formulae:

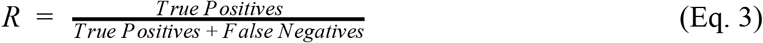

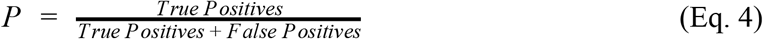

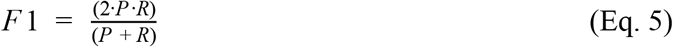

In this case, false negatives are criteria that were missed by SciScore but were labeled by a human curator, and false positives were incorrectly identified text labeled by SciScore.

We did not obtain an institutional review board approval to conduct this study as we did not utilize any human or animal subjects, making this study exempt.

## Results

Of the 1,578,964 articles analyzed by SciScore, 197,892 articles were considered not applicable. Research was considered not applicable if no scoring criteria were found resulting in a SciScore of 0. In total, 1,381,072 papers were scored, giving a score rate of 87.5% for articles with accessible methods sections. To determine the quality of SciScore annotations beyond the training set, we used a sampling of 250 papers randomly selected from the scored papers (SciScore > 0) and 250 papers randomly selected from the “not applicable” papers (SciScore = 0). Every methods section was read, and the curator noted the presence or absence of each entity type used in our analysis (entity types shown in Table 1). As previously mentioned, some entity types (IRB and consent; cell line authentication and contamination) were combined. In order to maintain consistency throughout our analysis, we counted the presence of one of these entity types as sufficient for both. Of these entity types though, all can be considered conditional and are therefore not entirely independent; e.g., studies that require IRB approval usually require a statement of consent; studies using cell lines normally require both an authentication statement and a contamination statement. Because of this, we feel that it is not unreasonable to group these criteria together in these instances.

For the 250 scored papers, the curator-SciScore agreement rate for each entity type is shown in Table 2. These 250 papers were scored by a curator for each of the criteria listed in Table 2. In every case, the entity type had an agreement rate above 80%; most were over 90%. We assumed that if both the curator and SciScore agreed about the presence of an entity type in the paper, then the answer was correct and we did not look more deeply into these data. We reported papers with disagreement as either false positives or false negatives with the assumption that the curator is always correct. False negatives were defined as cases where the classifier incorrectly noted an entity type as absent when it was in fact present. Inversely, false positives were defined as cases when the classifier incorrectly noted an entity type as present when it was missing. The false negative values and false positive values for each entity type are listed in Table 2. For key resources, the overall agreement should represent additive probability for instances where multiple resources were mentioned. In all cases, the agreement rate was measured above the raw classifier F1 rate except for software tools, which had an agreement rate that was lower than expected based on our previous training data (Table 1).

Of the 250 “not applicable” papers, 81.2% were found to have been correctly scored (n = 203). Of these 203 papers, 5 were found to be using supplementary methods sections, so a human might be able to look at these, but these sections are invisible to our algorithm, so we did not consider these a miss; 6 had their experimental procedures broken up across different sections of their papers, while 6 did not contain a clear methods sections at all. 47, or 18.8%, of the “not applicable” papers were found to have been incorrectly scored, that is, they were within scope, but the algorithm did not detect any relevant entity. Of these 47 incorrectly scored papers, 45 were found to contain at least one software tool that was not detected by SciScore. This was by far the most missed entity in this set of papers. Blinding and sex as a biological variable were each missed by SciScore in 3 papers, while IRB/Consent, IACUC, blinding, and organism entity types were each found to only have been missed in one paper. These values all fall in line with what was expected based on our calculated rates for false negatives (shown through the recall rate in Table 1). The relatively low agreement rate for software tools seems reasonable as new software tools are often created with a specific use in mind and, as a result, are sometimes only used a handful of times. Because of this, there is a high number of uncommon software tools with which SciScore has very little tool specific training data. This leads to a higher rate of false negatives for those types of software. However, this issue only impacts uncommonly used or recently created software. As a result of these analyses, we did not seek to tune parameters further for SciScore.

### Reproducibility criteria over time

In table 3, scored PMC-OAI data is grouped by journal and year showing the average SciScore, RTI, the proportions of papers addressing specific rigor criteria, and the proportions of uniquely identifiable resources. In total, only 8 papers received the maximum score of 10. Summary data is presented to preserve author anonymity.

[Table 3 Supplemental Data: All Journals scored by year]

For reference, the average SciScores, standard deviations, proportions of papers addressing rigor criteria, and proportions of identifiable key resources across all papers (SciScore > 0) are shown in the supplemental data file for Figure 1. This data is grouped by year (1997 - 2019).

Between 1997 and 2019, the average, annual SciScore has more than doubled from 2.0 ± 0.9 to 4.2 ± 1.7 (Fig 1.A). This increase in SciScore coincides with increased levels of both rigor criteria inclusion and key resource identifiability. For rigor criteria inclusion, adherence levels largely increased for the following criteria: sex (21.6% to 37.0%) and randomization of group selection (9.8% to 30.1%). Levels of inclusion of statements about blinding (2.9% to 8.6%) and power analysis (2.2% to 9.9%) increased, but remained relatively low (Fig 1.B). For key resource identifiability, antibodies (11.6% to 43.3%) and software tools (42.1% to 86.7%) were increasingly found to be uniquely identifiable in the methods section, while organisms (21.1% to 22.0%) and cell lines (36.8% to 39.3%) remained at low levels of identifiability (Fig 1.C).

**Figure 1:**
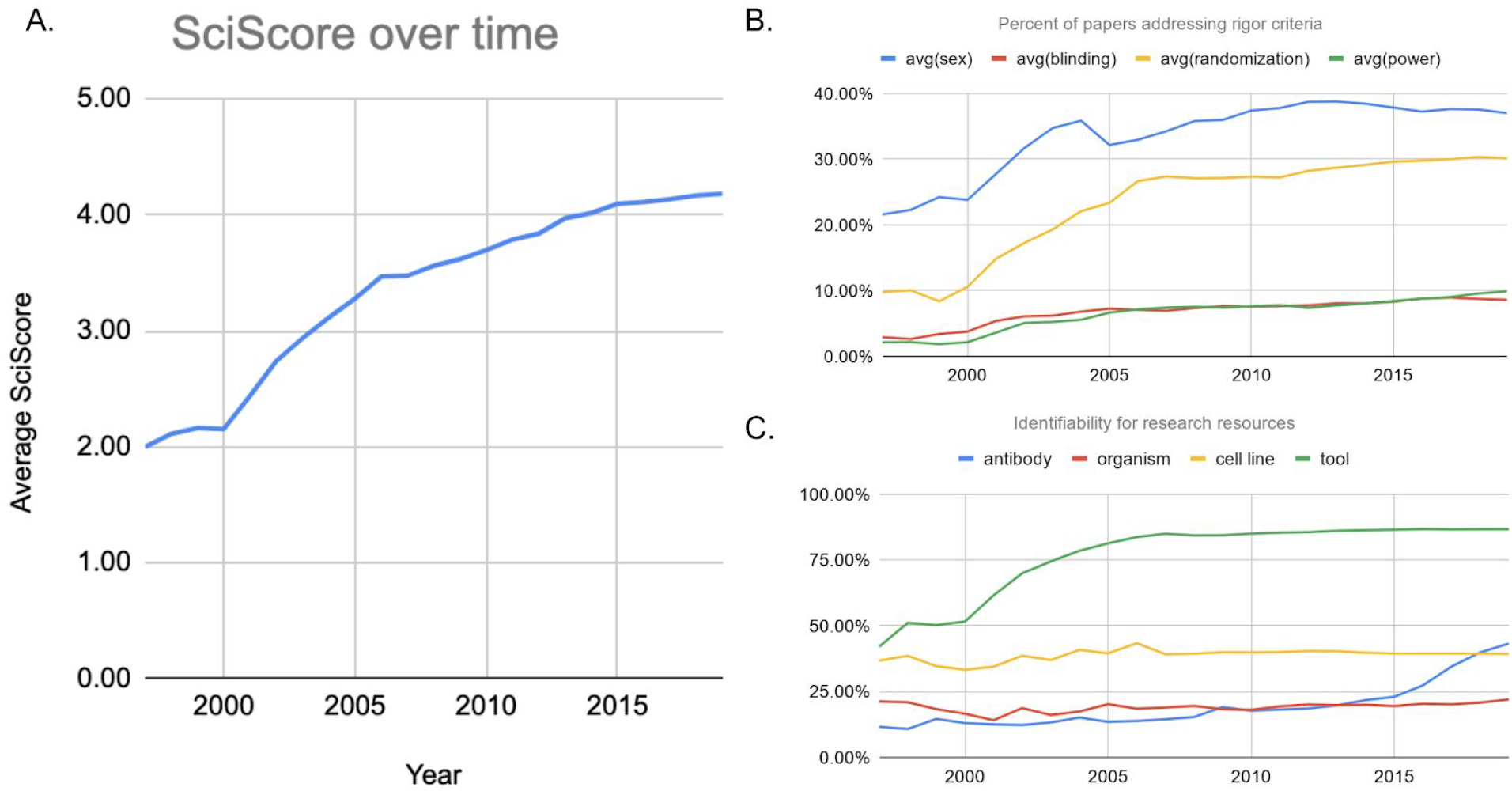
Overall SciScores and their breakdown shown between 1997 and 2019. (A) Average SciScore of the scored dataset representative of the biomedical corpus showing a relatively steady increase over time. (B) Percentage of papers mentioning the use of sex, blinding, randomization of subjects, and power analysis. Sex and randomization have increased significantly, while blinding and power analysis have increased, but are still at relatively low rates. (C) Percentages of key resources (antibodies, organisms, cell lines, and software tools) that are considered uniquely identifiable. Rates of software tools and antibodies have increased, while organisms and cell lines have remained relatively stagnant. Data underlying these graphs are available in the supplemental data file for figure 1.

To serve as a control, we analyzed only the most relevant papers to determine if our overall analysis accurately reflects more granular subsets. Papers containing IACUC statements in 2018, the most recent full year, comprised about a quarter of the total PMC-OAI papers scored (51,312 of 208,963). These animal papers showed the following rates of rigor criteria: sex 55.82% (compared to 37.54% in the total set), blinding 12.33% (8.74%), randomization 36.26% (30.30%), and power 7.34% (9.57%). Of 62,652 organisms detected in the total literature, 51,134 were represented in the subset. Identifiability of organisms was 21.71% vs 20.81% in the total set. These numbers suggest a trend in that the vertebrate animal subset of the literature is somewhat better than the total literature especially when looking at sentences describing the sex of the animal, group selection criteria and blinding, it remains far from ideal.

Antibody identification, in particular, has made considerable improvements going from the least identifiable key resource to the second most identifiable one in just the last few years, although antibody identification still remains under 50% overall. Some journals have made significant changes, leading to a more dramatic improvement compared to others (Fig 2). For example, Cell, a STAR^20^ methods compliant journal, improved their antibody identifiability rates from 11.1% to 96.7% from 2014 to 2019. eLife, a participant in the RRID Initiative, increased their antibody identifiability rates from 27.2% to 89.6% from 2014 to 2019. On the other hand, Oncotarget (21.6% to 36.4%) and PloS One (22.7% to 32.2%) have each improved, but their absolute rates remain relatively low, with each falling below the overall average during that time frame (21.8% in 2014; 43.3% in 2019). For this analysis, we only included journals that had more than 10 papers, containing at least one antibody, scored (SciScore > 0) in a given year.

**Figure 2:**
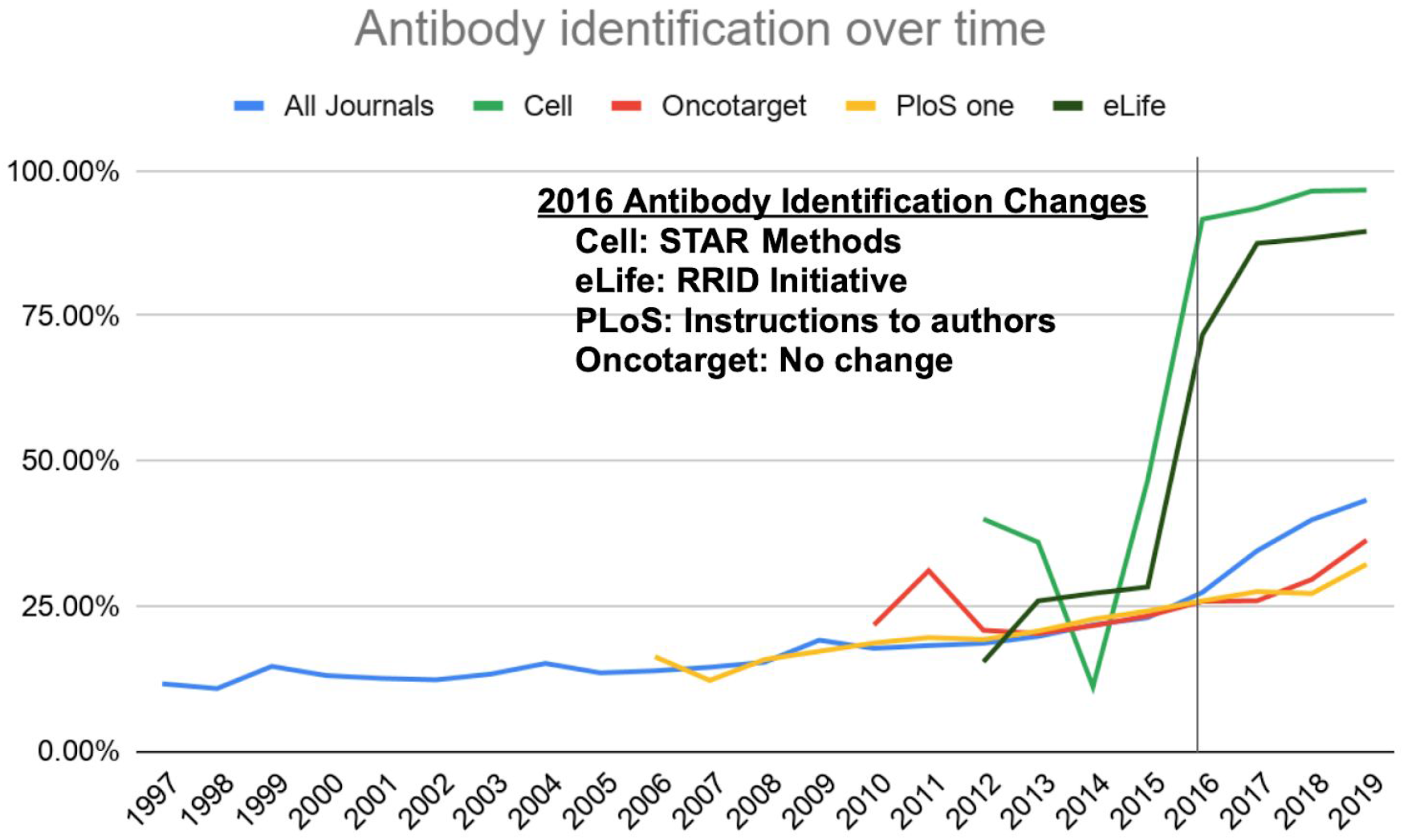
Percentage of antibodies that are able to be uniquely identified shown by journal with the overall trend across the biomedical literature shown in blue. A significant improvement can be seen starting in 2016 for Cell and eLife when STAR methods formatting and RRIDs were first implemented in their respective journals contributing to a noticeable improvement in antibody identifiability for the entire biomedical literature. Data underlying this graph are available in the supplemental data file for figure 2.

Table 5 shows the top 15 journals with the highest antibody identification rates for 2019 along with the number of antibodies detected in each. Seven (46.7%, Cell Stem Cell, Immunity, Cell, Molecular Cell, Developmental Cell, Cell Metabolism, Current Biology, and Cell Reports) have implemented the STAR methods reagent reporting format. Fourteen (93.3%, see Table 5) participated in the RRID Initiative and continue to enforce the use of RRIDs as of 2019. Therefore, these two drivers (STAR methods implementation and RRID Initiative participation) appear to have meaningfully contributed to improving the rate of identifiability in a majority of the best antibody identifying journals.

**Table 5:**
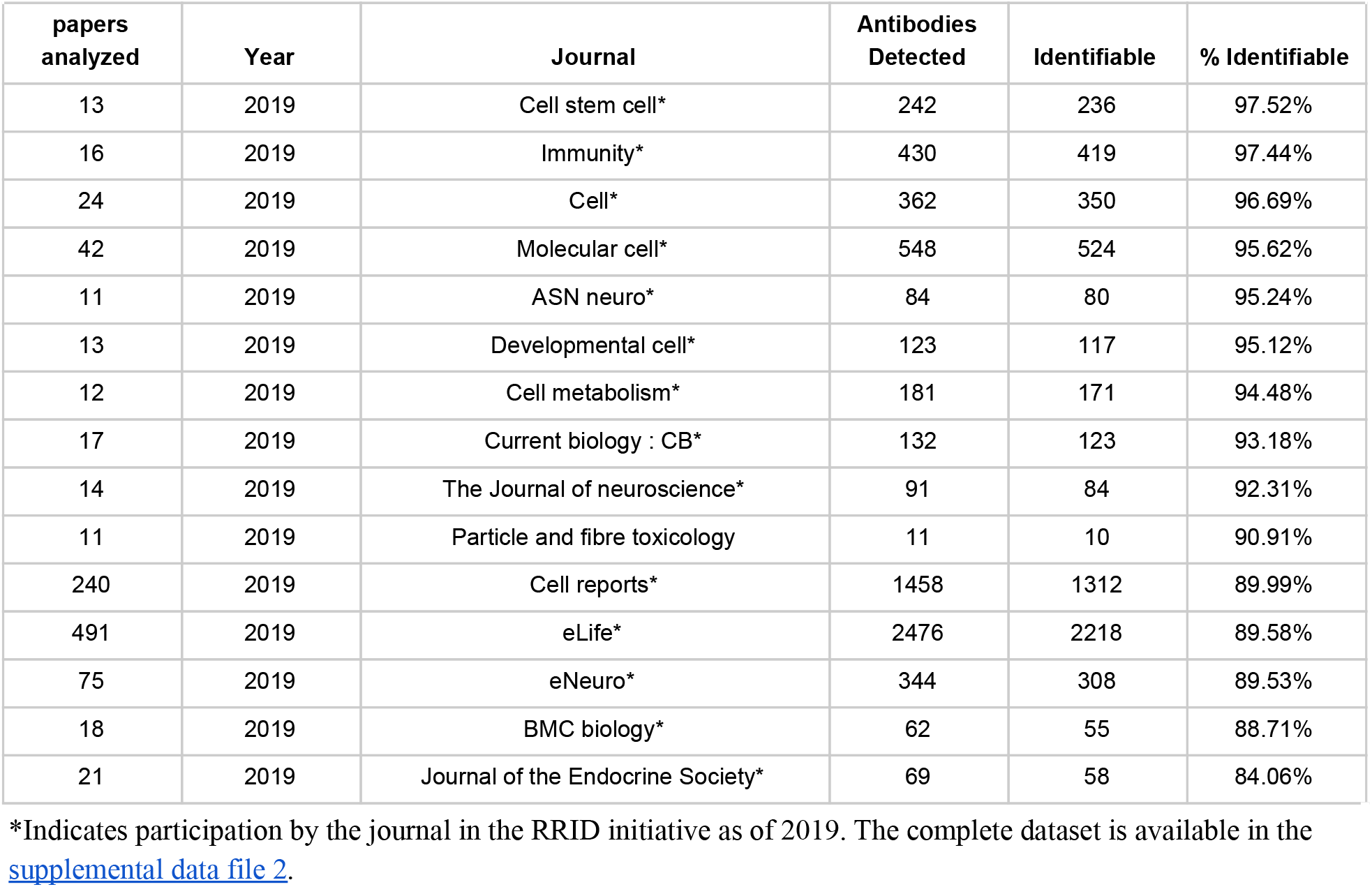
Top 15 journals sorted by percent of antibodies that were identifiable in 2019. For this analysis, there were 682 journals in which more than 10 antibody containing articles were accessible in our dataset.

All cell lines should be authenticated according to the international cell line authentication committee (ICLAC) guidelines because cell lines often become contaminated during experiments.^21^ Authentication of cell lines is usually accomplished by short tandem repeat (STR) profiling. This procedure is recommended at the outset of the experiment, at the conclusion of the experiment, and at a random time during the experiment. If this important control is completed, it should be stated in the manuscript. Similarly, authors should also test whether mycoplasma has contaminated their cell lines. For our purposes, we treated checking for mycoplasma contamination and authentication assessment like STR profiling as evidence that authors checked at least some aspect of cell line authenticity. Table 6 shows the journals that have the highest rates of authentication or contamination and the identifiability of cell lines in those journals. The percent of authentication is calculated as the percent of papers that contain a contamination or authentication statement is detected where at least one cell line is found.

**Table 6:** Top 15 journals sorted by percent of cell line authentication (authentication or contamination) that were identifiable in 2019. There were 2,280 journals in which more than 180,316 articles and more than 388,337 cell lines were accessible in our dataset. The complete dataset is available in the supplemental data file 3.

| **papers analyzed** | **Year** | **Title** | **Cell lines found** | **% Identifiable** | **% Authentication** |
| --- | --- | --- | --- | --- | --- |
| 278 | 2019 | eLife | 849 | 54.77% | 70.86% |
| 11 | 2019 | Nature microbiology | 39 | 33.33% | 63.64% |
| 83 | 2019 | Oncogene | 302 | 41.06% | 54.22% |
| 23 | 2019 | Journal of cell science | 87 | 31.03% | 52.17% |
| 27 | 2019 | Nature | 85 | 47.06% | 51.85% |
| 33 | 2019 | Oncogenesis | 95 | 35.79% | 51.52% |
| 14 | 2019 | Breast cancer research | 86 | 34.88% | 42.86% |
| 29 | 2019 | EMBO molecular medicine | 65 | 38.46% | 41.38% |
| 17 | 2019 | Disease models & mech. | 23 | 43.48% | 41.18% |
| 17 | 2019 | EMBO reports | 42 | 35.71% | 41.18% |
| 18 | 2019 | Cell | 67 | 35.82% | 38.89% |
| 178 | 2019 | BMC cancer | 485 | 40.21% | 38.20% |
| 37 | 2019 | Molecular cell | 157 | 36.94% | 37.84% |
| 14 | 2019 | Ther. adv. in me. oncology | 48 | 43.75% | 35.71% |
| 946 | 2019 | Nature communications | 2601 | 38.06% | 34.14% |

Checklists may assist authors in finding aspects of their manuscript that were not addressed, but until now it has been very difficult to determine if these checklists are effective. Most studies that addressed this issue looked at a relatively limited sample of journal articles.^12^ We consider below a use case, in which the implementation of a checklist system appeared to be effective in improving the number of rigor criteria addressed by authors. In 2013 to 2014, Nature made a significant push with authors to address rigor criteria. We plotted the average SciScore along with its components over this period (Fig 3) and found that the average score rose by nearly 2 points over just a few years. This is based largely on a concomitant rise of authors addressing blinding, randomization and sex of subjects. To a smaller degree, antibodies became more identifiable and power analysis was described in a larger proportion of papers. In stark contrast, the Proceedings of the National Academy of Sciences of the United States of America (PNAS), which put out several reports advocating for the need for increased rigor,^22^ showed no change in composite score: 3.33 in 2015 to 3.42 in 2019.

**Figure 3:**
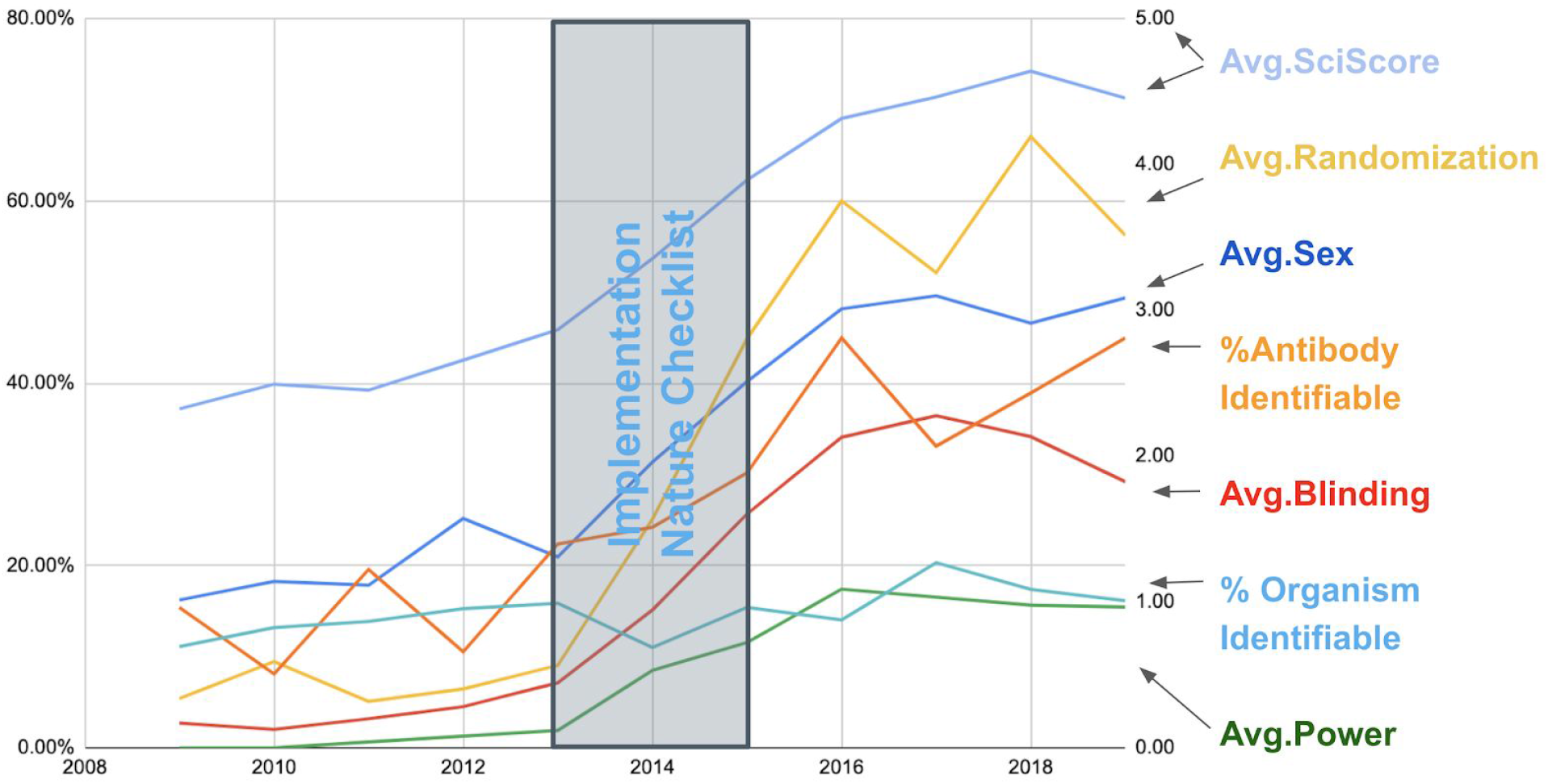
Analysis of rigor criteria for the journal Nature. The left axis represents the percentage of papers that fulfill a particular criterion. The right axis represents the average SciScore. The figure shows that during and after the implementation of the Nature checklist, the average SciScore as well as all measures except for organism identifiability have improved markedly. While scores were increasing before the checklist implementation, the checklist appears to quickly boost numbers. Data underlying this graph are available in the supplemental data file for figure 3.

### A comparison of the Rigor and Transparency Index with the Journal Impact Factor

In total, we included data from 490 journals (totaling 243,543 articles) for the JIF vs. Journal Rigor and Transparency (average SciScore) comparisons. The comparison between the raw JIFs and the Rigor and Transparency Index (Fig 4.A) showed a slight negative relationship, however, the correlation coefficient (R_s_ = −0.1102) suggests that this is not a significant relationship. Similarly, the JIF percentile vs. SciScore percentile relationship showed no significant correlation (R_s_ = −0.1541; Fig 4.B).

**Figure 4:**
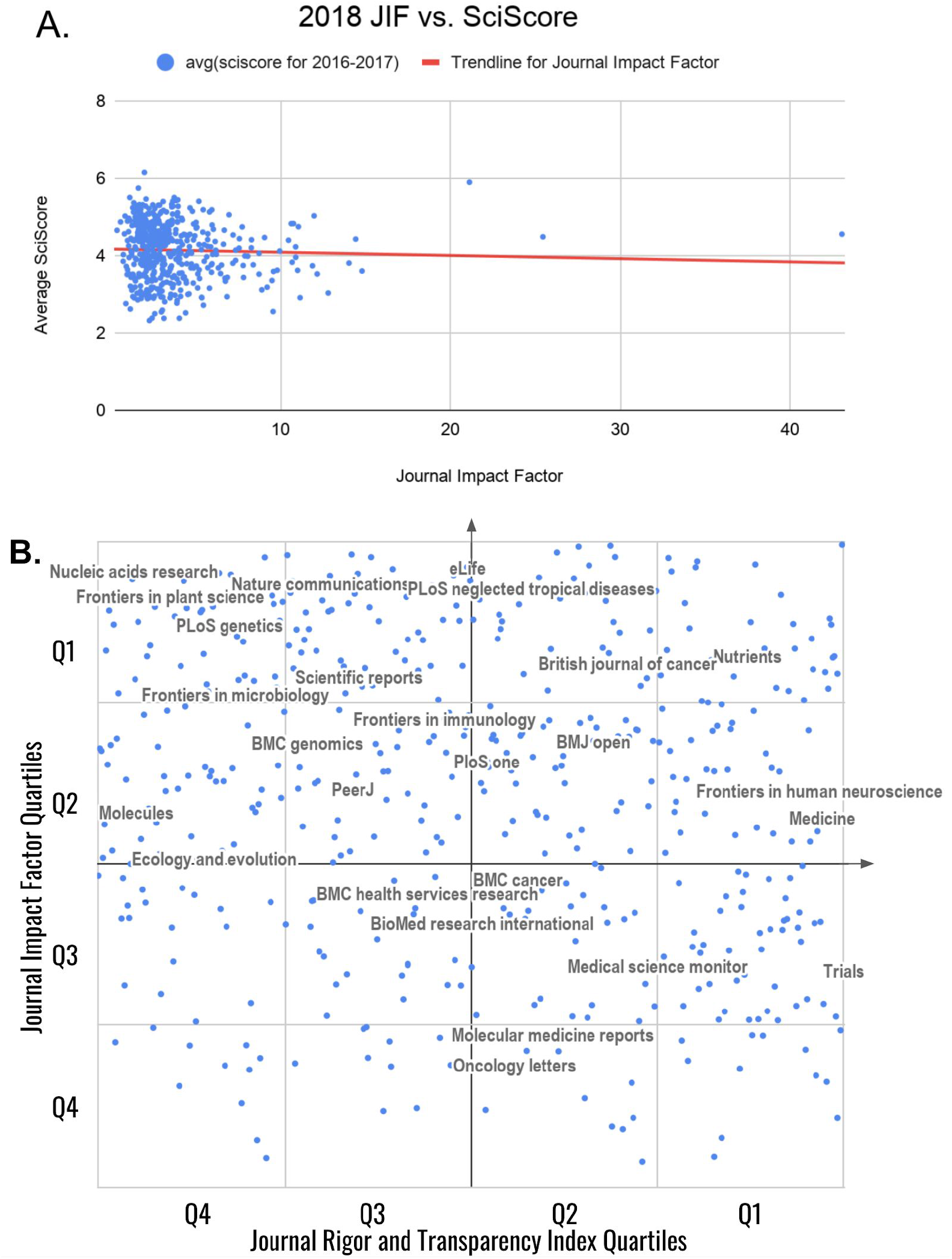
Average journal SciScore between 2016-2017 as a function of the journal impact factor for 2018 (data from published papers from 2016-2017). Data from 490 journals are shown in each graph. (A) A comparison between the raw JIFs and Rigor and Transparency Index is shown. The correlation coefficient is calculated using the formula for Spearman’s rank-order correlation (R_s_ = −0.1102253134). (B) A comparison between JIF percentiles and SciScore percentiles is shown. The axes are labeled with quartiles; top quartile is Q1. For presentation purposes only, using Google Sheets with journal names as centered data labels, we chose the top 45 journals by the number of articles included and then we removed labels that were overlapping until we were left with 25 labeled journals, shown above. All 490 journals, for which we had sufficient data in the open access literature to compare to the Journal Impact Factor, are presented in supplementary data file for figure 4. Correlation values were calculated using the formula for Spearman’s rank-order correlation, the line is not shown (R_s_ = −0.1541069707).

## Discussion

In this study, we introduce an automated tool, SciScore, that evaluates the materials and methods sections of a scientific paper for adherence to several key reporting guidelines introduced by funding agencies and journals over the past decade. Because the tool is automated, it provides us the opportunity to look at overall trends in adherence to these guidelines across the breadth of the scientific literature for the first time.

### Technical considerations

Currently, PMC-OAI currently represents only a fraction of the total biomedical literature and as a result must be considered a biased subsample. First, because this data consists of only full text accessible, open-access papers or copies of closed access papers supported by the NIH with licenses permissive for text mining, some journals may not be represented. Second, PMC was only launched in February 2000, meaning that papers in PMC will be on average more recent than the entire literature. Most of the data available through PMC is from the past 5 to 10 years, whereas PubMed contains a significant amount of older articles that date back 30 to 40+ years. Because of these concerns, we concede that PubMed, with 30.37 million articles as of November 27, 2019, is only partially represented in the portion of PubMed Central accessible for text mining. As a result of this differential, we cannot be certain that the text-mining accessible papers in PubMed Central are completely representative of the totality of biomedical literature. However, given that PubMed is our best guess at the totality of the biomedical literature, then it stands to reason that a sample of 5.2% of a population of this magnitude should be a reasonable representation of the total, especially in the more recent years. Journals that are not represented in this set of data are either those that are unavailable as open access, or unavailable under a text-mining allowed license.

We note that one of the most glaring omissions from our dataset is the journal Science. There are several reasons why this may be the case, the most likely being that the articles are not included in the PMC-OAI, a subset that is roughly half of the “free to read” set of papers in PubMed Central because of restrictive licenses. The other issue with Science articles specifically is that many of them do not contain a methods section that can be scored. Since Science’s format is highly abbreviated, the methods section tends to be pushed into the supplementary materials where it is likely to be formatted as an image (pdf file) rather than text. It is possible that this treatment of the methods section leads to not only invisibility from text mining algorithms, but also less attention from reviewers. To a text mining algorithm that expects text data formatted according to the journal article tag suite (JATS) standard, a pdf file labeled supplement is effectively invisible. One way to get around this is to attempt to score these papers manually. For the 18,000 papers available from Science in PubMed Central, a person who could score a paper every 5 minutes would need to work for roughly 1,500 hours or 187 days 24/7. As a point of reference, the algorithm was measured to score a paper in about 2 seconds, so the task would take 10 hours on a single machine (the 1.6M articles were processed in about 6 weeks on a single machine). The biomedical literature is being produced at a rate of about 2 million articles per year, a rate that long ago has exceeded the ability of any human to read, much less deeply understand the content. We expect that scientists will need a helping hand from some form of robot that can pre-digest some of this content, but to be effective, this robot will need access to the content. It would be a real shame if the flagship journals were not represented in this new paradigm.^23^

Several aspects of the SciScore classifiers had low F1 scores (Table 1), indicating that the algorithm had a relatively more difficult time in finding some types of entities. One example is power analysis, which had an F1 score of 64 with very high variability. This means that for this metric and also for cell line authentication (F1 = 54), the currently deployed algorithms are simply not all that accurate. The problem likely stems from the fact that our curators were only able to find 80 examples of power analysis statements and 150 examples of cell line authentication in the tens of thousands of papers that constituted our training dataset. Compared to the 17K statements involving antibodies, this number is very low. In the future, we plan to create an expanded data set to improve these numbers. However, the simple fact that curators could not easily find these statements in the literature also shows that these rigor criteria are the least used and most problematic.

### Analysis of reporting trends

Since the early 2000s, there have been multiple calls to improve scientific reporting and increase the specificity within methods sections because of irreproducible research.^24,25^ In 2007, Sena and colleagues used meta-analysis to assess the presence of various rigor criteria in the scientific literature about different diseases.^15^ While we are not able to exactly replicate those findings, our results can be compared. In their study, a human curator scored the presence of rigor criteria in 624 papers, a tremendous amount of human effort. These were broken down into disease groups, including stroke, multiple sclerosis and parkinson’s disease. In this set of the literature, authors addressed randomization of subjects into groups between 1 and 10% of the time. In our data, randomization is addressed in 8 to 27% of papers between 1997 and 2007, with a steady rise of this value over time. It is likely that Sena sampled from 2005 papers and before more frequently, making the range comparable. In the Sena paper, blinding was addressed 2 to 13% of the time depending on the disease area, while our data shows a range of 3 to 7% of papers where authors mentioned blinding. Sample size calculation was not detected in any study by Sena and our data shows a 2 to 7% detection rate of power calculations. Our data includes pre-clinical and clinical studies, while Sena’s study only included the former, making a direct comparison a little more tenuous. While reasonable people may argue that different techniques were used in performing these studies, including study selection and the criteria for inclusion, there is a striking similarity in this very cursory comparison, suggesting that the overwhelming majority of studies published in 1997-2007 did not address randomization, blinding and power analysis. This result is not entirely surprising given that these factors were specifically identified as leading to problems with reproducibility and were therefore targeted in the reporting guidelines that emerged after this period.

Since 2007, there has been a steady improvement in rigor inclusion and key resource identifiability rates across the literature. Between 1997 and 2019, the average score of biomedical research has more than doubled indicating an improvement in the transparent reporting of scientific research. However, it is difficult to assign causality. While the checklist implemented at Nature has clearly been well executed (see Fig 3), in general, guidelines and checklists have been shown to be relatively ineffective at improving the reporting tendencies of authors; because of this, we highly doubt these improvements are entirely due to the presence of checklists and guidelines.^12^ We do believe, however, that these guides provided authors with good focal points for where they should put forth effort in order to improve the reporting of their research, and while efforts such as the ARRIVE guidelines initially remained relatively unsuccessful in changing author behavior, there was eventual improvement (Fig 1). Given our current dataset, we can state that these reporting improvements appear to be occurring across biomedicine in general, suggesting that they may be due in part to an increase in awareness of the importance of reporting on good scientific practice.

While there are many causes contributing to the complex issue of scientific irreproducibility, none have been more vilified than the antibody.^26^ As one of the most prevalent tools in modern-day biological research, they represent an easy target raising the ire of disgruntled scientists as they are known in many cases to display a high level of variability between sources.^27^ These issues with antibodies, however, cannot be discovered in most papers because in most papers, even today, these reagents are not cited in a way that makes it easy to even understand which antibody has been used. Antibodies have long been one of least identified resources (Fig 1C).^10^ Comparing the Vasilevsky results to our current analysis, we found that antibodies are identifiable less often. For 2013, Vasilevsky found that ~45% of antibodies are identifiable, while our algorithm found it to be ~20% (in 2013). This discrepancy can likely be attributed to differences in criteria used, the exact papers analyzed, and the size of the sample. For each antibody, Vasilevsky looked in all vendor catalogs and searched for the name of that antibody, if the vendor search resulted in only one antibody, it was considered identifiable. This presents a bit of a best case scenario for antibody identification as similarly named antibodies may be added to a company’s inventory in the future or an antibody may simply have its name altered over time. Our algorithm relies on the presence of either a catalog number or a RRID for identifiability, which are far more stringent. While catalog numbers may still be quite imperfect for identifiability,^28^ they are nevertheless far more stable than a product’s name appearing in the vendor’s catalog. Additionally, RRIDs are significantly more stable than either a product’s name or its catalog number as they are meant to serve as a sort of unique product code, UPC, that transcends any of these superficial changes.

Data about software tools may be subject to more significant recency effects than other resources. This is partially because SciScore can detect that a word or phrase is the name of a software tool, but to be considered identifiable the tool must have accessible metadata. It is relatively more likely that tools in common use are more identifiable than tools that may have been used over a short period of time. Despite this, we still feel that SciScore has captured a majority of the mentioned software tools. This is because software tool mentions appear to follow the 80/20 rule where roughly 80% of the mentions are related to 20% of the tools.^29^

Our analysis of the antibody data clearly demonstrates that some journals, which enforce RRIDs, have dramatically higher rates of identifiability (>90%) than the average journal (~40%), see Table 5. Enforcing the use of RRIDs is not an effortless exercise; we understand from personal communications with the editors of the Journal of Neuroscience that authors are asked to identify their research resources 3 times during the publication process, which takes a substantial effort. The Cell Press family of journals is quite interesting because of the requirement for a STAR table, which makes antibody identification highly visible to journal staff and authors.^20^ These journals are the vanguard of rigor and should be celebrated, especially because it appears that they are not only moving their authors to change behavior, but that these changes in behavior are also spreading as evidenced by a fairly dramatic overall shift in identifiability since 2016. We do not know why this spread is occuring, but in seemingly unrelated journals that did not change policies with regard to antibody identification there are more well identified antibodies. Some of this could be explained as journals that enforce policy have high rejection rates and those authors end up in another journal with well identified antibodies. It may also be that authors have been frustrated for so long trying to track down antibodies that when they hear about a way to change the current practice, they embrace the change.

Through the use of a vastly different performance indicator than what is currently used (SciScore as opposed to JIF), we have created a method to score journals that is very different than the impact factor. The Rigor and Transparency Index lists journals with their composite scores and rates of inclusion for rigor adherence and resource identifiability. We choose to include JIF percentiles in our analysis because we thought it gave a more accurate measure of the “best” journals as JIFs are often only compared to journals in the same field due to variations in citation counts between different scientific branches.^30^ Using the average JIF percentile, we were able to account for these changes. We also feel that any impact associated with the FUTON or NAA (no abstract available) biases would be mitigated because a vast majority of the journals analyzed were at least partially open-access and all cases where abstracts were not available were universally excluded. In the end though, our analysis indicates that there is no correlation between a journal’s impact factor and the Rigor and Transparency Index.

Researchers have pointed out various problems with measuring journals based on the JIF.^31,32^ Many of these arguments are valid in that they point to this single number as an “outdated artifact” that improperly impacts how we view research. The most important underlying problem with the JIF, in our opinion, is that it measures popularity (number of citations) and not the quality of the work.

The Rigor and Transparency Index differs from the JIF in that it is based on known problem areas linked to the inability to reproduce a study. While the composite number for any given study is likely nearly meaningless (an 8 is not demonstrably better than a 7, for example), it is very difficult to argue that reagents used in a study should not be referenced in such a way as to easily identify them. It is also true that all means of reducing investigator bias, such as blinding, are not possible in all experimental designs, especially during the conduct of certain experiments. However, it is difficult to argue that addressing investigator bias is a waste of time; indeed, investigators surveyed by Nature overwhelmingly state that the checklist which covers bias was helpful to their reporting of research.^33^ Investigator bias can creep into any scientific discipline and has been shown to artificially inflate effect size in stroke research,^34^ but these effects have been well understood since the 1960’s,^35^ and have informed the practice of clinical trials. While at the current state of the art it is nearly impossible to determine if authors are addressing rigor criteria appropriately in a particular study, the fact that most authors largely ignore these does mean that investigator bias is not “on the radar” of many researchers as they report on findings. In more general terms, we believe that research that completely and transparently reports its reagents and methods, is likely to be much better than research that does not. We therefore argue that a study that scores 8 or 9, which will necessarily address investigator bias and uniquely identify most resources, is better than a study that does not address these and scores 2 or 3.

The creation of the Rigor and Transparency Index provides both a short hand for how a journal is doing, and a much more detailed picture of the current state of rigor and transparency practices. It can point each journal to significant problem areas that are addressable in future publications. It also provides journals and funders the ability to monitor the impact of their policies regarding rigor and reproducibility. The RTI can bring attention to the importance of sound science practices.

## Acknowledgements

We thank the following individuals for their contribution to this work: Gabrielle Pine, who provided substantial insight with the writing of this manuscript; Nathan Anderson, Edyta Vieth, and Ethan Badger, who helped to create many of the datasets and validation series. We also thank Dr Anthony C. Gamst for checking our statistical assumptions. Moreover, we thank the many journal editors and editorial staff who continually work to improve the research integrity by augmenting their workflow to include RRIDs. NIH This work was funded by: OD024432, DA039832, DK097771, MH119094, and HHSN276201700124P.

## Notes

#### Summary of Updates

We forgot the Acknowledgements in our original submission.

https://docs.google.com/spreadsheets/d/1gAtik_zqV1iIi7GFYrIwkoG8cq7gPJRrjthCz0MZpfs/edit?usp=sharing

